# Retrieval of a well-established skill is resistant to distraction: evidence from an implicit probabilistic sequence learning task

**DOI:** 10.1101/849729

**Authors:** Teodóra Vékony, Lilla Török, Felipe Pedraza, Kate Schipper, Claire Plèche, László Tóth, Karolina Janacsek, Dezso Nemeth

**Affiliations:** Department of Neurology, University of Szeged, Szeged, Hungary; Department of Psychology and Sport Psychology, University of Physical Education, Budapest, Hungary; Université Lumière - Lyon 2, Lyon, France; Lyon Neuroscience Research Center (CRNL), INSERM, CNRS, Université Claude Bernard Lyon 1, Bron, France; Institute of Psychology, ELTE Eötvös Loránd University, Budapest, Hungary; Brain, Memory and Language Research Group, Institute of Cognitive Neuroscience and Psychology, Research Centre for Natural Sciences, Hungarian Academy of Sciences, Budapest, Hungary; School of Human Sciences, Faculty of Education, Health and Human Sciences, University of Greenwich, London, United Kingdom

**Keywords:** implicit sequence learning, statistical learning, memory retrieval, dual-task, divided attention

## Abstract

The characteristics of acquiring new sequence information under dual-task situations have been extensively studied so far. Such a concurrent task has often been found to affect performance. In real life, however, we mostly perform a secondary task when the primary one is already well-acquired. The effect of a secondary task on the ability to retrieve well-established sequence representations remains elusive. The present study investigates whether accessing a well-acquired probabilistic sequence knowledge is affected by a concurrent task. Participants acquired non-adjacent regularities in a perceptual-motor implicit probabilistic sequence learning task. After a 24-hour offline period, participants were tested on the same sequence learning task under dual-task or single-task conditions. Here we show that although the secondary task significantly prolonged the overall reaction times in the primary task, the access to the previously learned probabilistic representations remained intact. Our results highlight the importance of studying the dual-task effect not only in the learning phase but also during memory access.

## Introduction

Sequence learning is a fundamental function of the brain, which underlies the acquisition of motor, cognitive and social skills (Kaufman et al., 2010; Lieberman, 2000; Nemeth et al., 2011; Romano Bergstrom, Howard, & Howard, 2012). These skill-based actions, such as driving a car or playing sports, usually become more automatic with extensive practice. In everyday life, these skills are generally not performed in isolation but simultaneously with other actions. Therefore, the effect of a secondary task on implicit sequence learning performance has been studied extensively in the last few decades. Depending on the characteristics of the primary and the secondary task, evidence for impaired (e.g. Franklin, Smallwood, Zedelius, Broadway, & Schooler, 2016; Hemond, Brown, & Robertson, 2010; Jiménez & Vázquez, 2005; Nissen & Bullemer, 1987; Röttger, Haider, Zhao, & Gaschler, 2017; Shanks, Rowland, & Ranger, 2005; Wierzchon, Gaillard, Asanowicz, & Cleeremans, 2012), intact (e.g. Jiménez & Méndez, 1999; Jiménez & Vázquez, 2005; Nemeth et al., 2011; Röttger et al., 2017; Rowland & Shanks, 2006) or even improved performance (Hemond et al., 2010) was found during the acquisition of implicit sequence knowledge. However, the effect of a secondary task on the *retrieval* of a complex, well-established skill is rarely studied. In everyday life, we mostly perform a secondary task when the primary one is well-learned. For example, when we are learning how to drive, our entire attention is on the primary task, and we refrain from other concurrent activates, such as chatting with our co-pilot. However, after mastering this skill, we easily engage in conversations during the primary (driving) task. Therefore, answering the question of whether our performance is affected in such cases is crucial in understanding the effects of a secondary task on real-life performances. Here we present a study where we test the effect of a secondary task *on the retrieval* of well-established perceptual-motor implicit sequence knowledge.

What do we know about the effect of a dual-task condition on the retrieval of a learned sequence? Sequence learning is typically measured by asking participants to respond to a series of stimuli that follow a predetermined sequence order (Nissen & Bullemer, 1987). By contrasting the performance to stimuli presented in a random order, the degree of sequence knowledge can be measured. Using short single-task practice periods and immediate retrieval, early studies have found impaired (Curran & Keele, 1993) or intact retrieval of sequence knowledge under dual-task conditions (Frensch, Lin, & Buchner, 1998; Shanks & Channon, 2002). The latter results were often interpreted as the secondary task affecting only the performance measured at the time of testing (i.e. the expression of knowledge), but not the underlying representations (Frensch et al., 1998). Cohen and Poldrack (2008) have investigated the effect of extended dual-task practice, and have demonstrated that the impairing effect of the dual-task decreases with extended practice on a serial reaction time task. These results raise the possibility that sequence learning knowledge remains intact after gaining experience on the primary task. These studies (along with most of the previous experiments testing the effect of dual-tasking on the initial learning) used fixed (deterministic) sequence learning tasks that might be less implicit than probabilistic tasks (Cohen & Poldrack, 2008; Frensch et al., 1998; Kóbor, Janacsek, Takács, & Nemeth, 2017; Robertson & Cohen, 2006). In a typical probabilistic sequence learning task, certain sequence elements occur with a higher probability than others, and participants learn to answer faster and more accurately for the more probable elements compared to the less probable ones (Howard & Howard, 1997). The probabilistic learning results in a robust knowledge (Kóbor et al., 2017), and it is more likely to find intact sequence learning when a concurrent secondary task is performed in the initial learning phase (Jiménez & Méndez, 1999, 2001); however, others did claim detrimental effects (Shanks et al., 2005). An early study of Schvaneveldt and Gomez (1998) have found that probabilistic sequence knowledge learned without a secondary task cannot be transferred to a dual-task condition. On the contrary, transfer from dual-task learning to single-task performance did occur, and the authors concluded that the impaired performance was owing to problems in the expression of knowledge, and not to the impaired learning itself. However, what do we know about the effects of a secondary task on the retrieval of an already acquired knowledge? There is a long history of studying the dual-task effects on automatic behaviors, mostly using non-complex, choice-response tasks. Most of the results support the claim that an already automatized behavior is resistant to concurrently performing a secondary task (e.g. Logan, 1979) and that the dual-task cost on *general skill learning* (i.e. on the increasing speed due to practice independently from the sequence structure) tend to decrease with extended practice (e.g. Brown & Bennett, 2002; Hazeltine, Teague, & Ivry, 2002; Ruthruff, Johnston, & Van Selst, 2001; Ruthruff, Van Selst, Johnston, & Remington, 2006; Van Selst, Ruthruff, & Johnston, 1999). Nevertheless, to the best of our knowledge, no study has compared directly the accessibility of more complex, well-acquired probabilistic sequence knowledge (learned without a secondary task) between single and dual-task testing conditions.

In the present study, we aimed to investigate the effect of a concurrent secondary task on the retrieval of implicit probabilistic sequence knowledge. So far, studies have investigated the effect of single-task practice on immediate retrieval of the sequence knowledge. However, we do not know whether a newly-introduced secondary task might disrupt the retrieval of a well-acquired sequence knowledge, although it resembles how we pursue dual-task situations in everyday life. Therefore, we investigated the effect of a secondary task on the retrieval of a well-learned probabilistic sequential knowledge after extended practice, a 24-hour offline period, and a reactivation phase (Figure 1). First, we trained all participants on a perceptual-motor probabilistic sequence learning task in a single-task condition. After a 24-hour offline period, participants were tested again; however, at this stage, they performed the task with or without a stimulus-counting secondary task. To control for the possibility that the potential differences emerge due to the difficulties in the expression of knowledge (Jiménez & Méndez, 1999; Vékony et al., 2019), we inserted single-task blocks into the retrieval phase of the dual-task group. Previous diverse findings did not allow us to make clear predictions about the results; however, three outcomes are conceivable for our study. First, as the majority of previous literature reported deteriorating effects of a secondary task on sequence learning abilities (Franklin et al., 2016; Hemond et al., 2010; Jiménez & Vázquez, 2005; Röttger et al., 2017; Shanks et al., 2005; Wierzchon et al., 2012) explained by numerous theories such as the suppression hypothesis (Frensch et al., 1998; Frensch, Wenke, & Rünger, 1999), task integration (Schmidtke & Heuer, 1997), organizational hypothesis (Stadler, 1995) or the response selection hypothesis (Schumacher & Schwarb, 2009), it is possible that the sequence knowledge retrieval will be impaired under dual-tasking. The second potential outcome is that performing a concurrent task would not influence the sequence knowledge retrieval. This would suggest that the process of sequence knowledge retrieval and the stimulus-counting secondary task can be independent from each other, the same way as performance is automatized and resistant to distraction on simple choice-response tasks (e.g. Brown & Bennett, 2002; Hazeltine et al., 2002; Logan, 1979; Ruthruff et al., 2001, 2006; Van Selst et al., 1999). The third possibility is that the secondary task would improve the retrieval of the sequence learning task that would fit in well with the hypothesis of competition between control functions and sequence learning abilities (Ambrus et al., 2019; Filoteo, Lauritzen, & Maddox, 2010; Janacsek, Fiser, & Nemeth, 2012; Nemeth, Janacsek, Polner, & Kovacs, 2013; Tóth et al., 2017; Virag et al., 2015).

**Figure 1.**
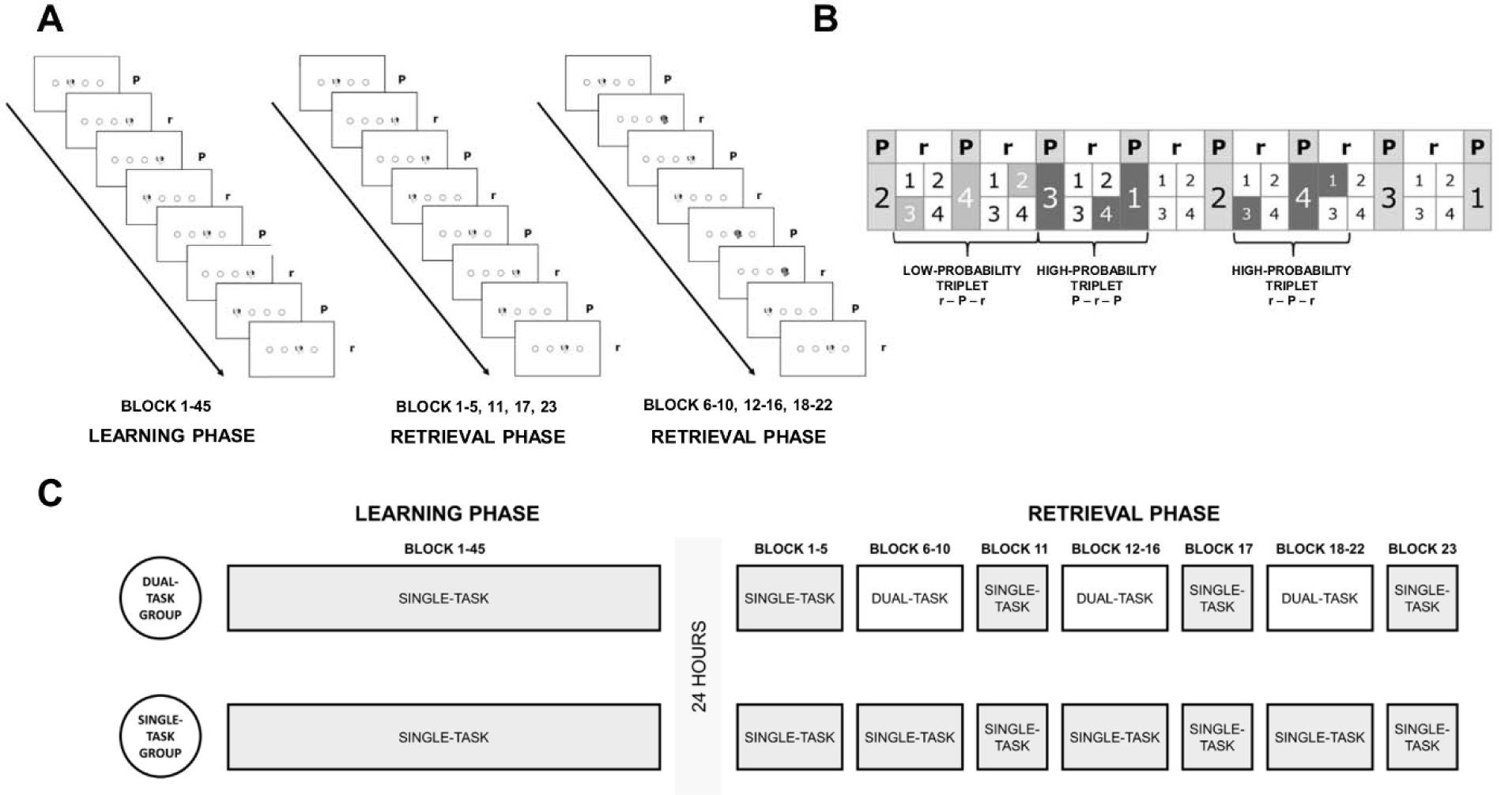
The structure of the ASRT and the experiment. (A) A dog’s head appeared on one of the possible positions. The participants’ task was to press the corresponding button as fast and as accurately as possible. In the Learning Phase, only black-and-white stimuli appeared. During the Retrieval Phase, in Blocks 6-10, 12-16 and 18-22, sometimes yellow stimuli appeared on the screen, which the participants in the Dual-Task Group had to count, while the participants in the Single-Task Group were told to ignore. (B) High- and low-probability triplets. Some runs of consecutive trials (triplets) occurred with a higher probability than other elements. Every stimulus was defined as the third element of a high- or a low-probability triplet, based on the two preceding trials. High-probability triplets can be formed by two patterns and one random element and – although more rarely-by two random and one pattern elements. (C) Experimental design. In the Learning Phase, both groups of participants practiced single-task ASRT throughout 45 blocks. After 24 hours of rest, in the Retrieval Phase, participants completed 5 more blocks of single-task ASRT (reactivation phase), and after that, 18 blocks of ASRT with or without dual-tasking (Dual Task Group and Single Task Group, respectively). However, 3 of them were probe blocks for both groups (Block 11, 17 and 23).

## Methods

### Participants

The participants were selected from a pool of eighty-one participants. One participant who did not follow the dual-task instruction in the Retrieval Phase (see Procedure section in the Methods) was excluded from the pool. As similar initial performance is a crucial criterion in our design, we needed to make sure that the observed effects were not due to differences in the initial learning performance or the consolidation of the sequence knowledge. Therefore, the participants in the two groups were matched by learning performance the first session, by level of consolidation (change in learning performance between the end of the Learning Phase and the beginning of the Retrieval Phase), and by the performance at the beginning of Retrieval Phase (still without dual-task, see Procedure section in the Methods). The final analysis was carried out on 68 participants (N_Single-task group_ = 34, N_Dual-Task Group_ = 34). The participants were between 18 and 33 years of age (M_age_ = 22.91 years, SD_age_ = 3.48 years). The years of education ranged between 10 and 20 (M_years of education_ = 14.66 years, SD_years of education_ = 2.36 years). Handedness was measured by the Edinburgh Handedness Inventory (Oldfield, 1971). The Laterality Quotient (LQ) of the sample varied between −53.85 and 100 (−100 means complete left-handedness, 100 means complete right-handedness; M_LQ_ = 44.79, SD_LQ_ = 34.22). All participants had normal or corrected to normal vision, none had a history of any neurological and/or psychiatric disorder or reported taking any psychoactive medication at the time of the experiment. They performed in the normal range on the Counting Span task (Range_Counting Span_: 2.33-6, M_Counting Span_ = 3.81, SD_Counting Span_ = 0.89) and on the Digit Span task (Range_Counting Span_: 5-9, M_Counting Span_ = 6.28, SD_Counting Span_ = 1.13). The two groups did not differ in any of the demographic and cognitive characteristics (Table 1). The Research Ethics Committee of the Eötvös Loránd University (Budapest, Hungary) approved the study and it was conducted in accordance with the Declaration of Helsinki.

**Table 1.** Comparison of the two groups on age, years of education, handedness, short-term, and working memory performance

|  | *Dual-Task Group* | *Single-Task Group* | *Group comparison* |
| --- | --- | --- | --- |
|  | *M(SD)* | *M(SD)* | *(t-test results)* |
| Age (years) | 23.03 (3.42) | 22.79 (3.60) | $t(66) = -0.276, p = 0.783$ |
| Education (years) | 14.50 (2.27) | 14.82 (2.47) | $t(66) = 0.562, p = 0.576$ |
| Handedness (LQ) | 50.86 (35.07) | 38.73 (32.80) | $t(66) = -1.474, p = 0.145$ |
| Counting Span Score | 3.92 (0.96) | 3.69 (0.81) | $t(66) = -1.045, p = 0.300$ |
| Digit Span Score | 6.32 (1.15) | 6.24 (1.13) | $t(66) = -0.320, p = 0.750$ |

### Alternating Serial Reaction Time task

We used the Alternating Serial Reaction Time (ASRT) task to test the implicit sequence learning abilities of the participants (Howard & Howard, 1997; Nemeth et al., 2010). Four empty circles (300 pixels each) were presented continuously in front of a white background arranged horizontally in the middle of a computer screen. A target stimulus (a black and white drawing of a dog’s head, 300 pixels) was presented sequentially in one of the four empty circles. Participants were asked to respond with their middle and index fingers of both hands by pressing the button corresponding to the target position on a keyboard with four heightened keys (Z, C, B, and M on a QWERTY keyboard), each of the four keys corresponding to the circles in a horizontal arrangement. Participants were instructed to be as fast and as accurate as possible in the task (Figure 1A).

The serial order of the four possible locations (coded as 1, 2, 3, and 4) in which target stimuli could appear was determined by an eight-element probabilistic sequence. In this sequence, every first element appeared in the same order as the task progressed, while the second elements’ positions were randomly chosen out of the four possible locations (e.g., 2r4r3r1r; where r indicates a random position). Thus, some combinations of three consecutive trials (*triplets*) occur with a higher probability than other combinations. For example, 2_**4**, 4_**3**, 3_**1**, and 1_**2** (where ‘‘_” indicates any possible middle element of the triplet) would occur with high probability because the third element (bold numbers) could be derived from the sequence (or occasionally could be a random element as well). In contrast, 1_**3** or 4_**2** would occur with less probability because these triplets cannot be formed from two sequence elements and one random element. Therefore, the third element of a high-probability triplet is more predictable from the first element when compared to a low-probability triplet. There were 64 possible triplets in the task altogether. Sixteen of them were high-probability triplets, each of them occurring in approximately 4% of the trials, about five times more often than the low-probability triplets (0.8%). Overall, high-probability triplets occur with approximately 62.5% probability during the task, while the remaining 48 low-probability triplets only occur with 37.5% probability (Figure 1B). As the participants practice the ASRT task, their responses become faster and more accurate to the high-probability triplets compared to the low-probability ones, revealing statistical learning throughout the task (J. H. Howard & Howard, 1997; Kóbor et al., 2017; Song, Howard, & Howard, 2007; Unoka et al., 2017). Six different sequences were used across subjects, but the sequence for a given subject was identical throughout the entire experiment.

The ASRT task was completed in blocks, and each block contained 85 button presses (5 random elements at the beginning of the block, then the 8-element alternating sequence was repeated 10 times). At the beginning of each block, four empty circles were presented horizontally on the screen for 200 ms, and then the first target stimulus appeared. The target stimulus remained on the screen until the first correct response. Between blocks, the participants could rest a little and received feedback about their performance (average RTs and accuracy) on the screen. After five blocks, a longer (5 minutes) mandatory pause was inserted. We defined each trial as the third element of a high or low-probability triplet. Trills (e.g. 1-2-1) and repetitions (e.g. 1-1-1) were eliminated from the analysis because participants may show pre-existing response tendencies for these types of triplets (Howard et al., 2004; Janacsek, Borbély-Ipkovich, Nemeth, & Gonda, 2018; Takács et al., 2018; Unoka et al., 2017). The first button presses were also excluded from the analysis (first 5 random button presses, and the 6^th^ and the 7^th^, as they cannot be evaluated as the third element of a triplet). To facilitate data analysis and to increase the signal-to-noise ratio, every five blocks were collapsed into a larger unit of analysis.

### Procedure

The experiment consisted of two sessions (Figure 1C). In the Learning Phase, participants completed 45 blocks of ASRT (45 blocks divided into 9 units of analysis: Blocks 1-5, Blocks 6-10, Blocks 11-15, Blocks 16-20, Blocks 21-25, Blocks 26-30, Blocks 31-35, Blocks 36-40 and Blocks 41-45), which is long enough to acquire a stable statistical knowledge (e.g. Vekony et al., 2019). In the Retrieval Phase, which was held 24 hours after the Learning Phase, participants completed 23 blocks of ASRT with the same sequence that they learned the previous day (20 blocks divided into 4 units of analysis and 3 separate blocks: Blocks 1-5, Blocks 6-10, Block 11, Blocks 12-16, Block 17, Blocks 18-22, Block 23). In Blocks 1-5, the instructions were completely the same as in the previous day. This phase was included to strengthen the knowledge acquired on the previous day and to make sure that the two groups consolidated the knowledge to a similar level (see Supplementary Material). However, in Blocks 6-10, Blocks 12-16 and Blocks 18-22, a random number of stimuli (between 40-45 out of the 85 appearing stimuli in one block) was colored in yellow. The Dual-Task Group was instructed to count the number of yellow dogs throughout the block. After the completion of the block, the participants had to type in the number of yellow dogs they had counted. The yellow-colored stimuli appeared for the Single Task Group as well, however, they were instructed to carry on with the task without paying particular attention to the differently colored stimuli. The performance in the secondary task was evaluated by calculating the difference from the correct number of yellow stimuli divided by the total number of yellow stimuli for each unit of five blocks (thus, resulting in a percentage score of correctly counted yellow stimuli relative to the total number of yellow stimuli). Using this measure, we were able to control for the differences in the number of yellow stimuli across participants and for the fact that counting more stimuli is also considered as erroneous counting. If a participant’s overall differences score was over 15%, the participant was considered as not properly following the instructions and was excluded from the analysis (1 participant). Two probe blocks were inserted between the 3 dual-task phases (Block 11 and Block 17), and one other at the end of the session (Block 23). In these blocks, there were no yellow stimuli for the Dual-Task, nor for the Single Task Group.

Following the ASRT task, the Inclusion-Exclusion task was administered to check whether the participants developed conscious knowledge about the learned probabilistic regularities (see Supplementary Materials for details).

### Statistical analysis

To evaluate the performance in the ASRT task, we calculated the median reaction times (RTs) separately for the high- and low-probability triplets in every 5 blocks of the Learning Phase (45 blocks: Blocks 1-5, Blocks 6-10, Blocks 11-15, Blocks 16-20, Blocks 21-25, Blocks 26-30, Blocks 31-35, Blocks 36-40 and Blocks 41-45), in the Retrieval Phase (20 blocks: Blocks 1-5, Blocks 6-10, Blocks 12-16, Blocks 18-22), and in each 3 probe blocks in the Retrieval Phase (3 blocks: Block 11, Block 17 and Block 23). Only correct responses were included in the RT analysis. We chose to focus on the analysis of RTs, as previous similar ASRT studies have observed ceiling effects in accuracy (e.g. Janacsek et al., 2018).

To test the effects of the secondary task, we compared (1) the performance *while* the Dual-Task Group performed the dual-task (test) blocks, and (2) the performance *between* the dual-task phases (probe blocks). For the former, we performed a mixed-design ANOVA with Triplet (high-vs. low-probability) and Blocks (Blocks 6-10 vs. Blocks 12-16 vs. Blocks 18-22 in the Retrieval Phase) as within-subject factors, and with Group (Dual-Task Group vs. Single-Task Group) as a between-subject factor. For the latter, we collapsed the performance of the three blocks together, as one block does not contain enough trials to reveal between-block effects. Then we ran a mixed-design ANOVA with Triplet (high-vs. low-probability) as a within-subject factor, and with Group (Dual-Task Group vs. Single-Task Group) as a between-subject factor. Moreover, we directly compared the *learning scores* (RTs for low-probability triplets – RTs for high-probability triplets) between the test and probe blocks in the two groups to test if the sequence knowledge is different during the two phases. To this aim, we performed a mixed-designed ANOVA with the Block Type factor (test blocks vs. probe blocks) as a within-subject factor and the Group (Dual-Task Group vs. Single-Task Group) as a between-subject factor

As the dual-tasking caused major differences in median RTs between the two groups in the Dual-Task Blocks, we wanted to make sure that the results found about the sequence knowledge were not due to the changes in the overall speed (i.e., because of the effect of the dual-task on the *general skill learning*). To this end, we performed an additional analysis of the data with standardized scores. The standardized RT scores were calculated by dividing the learning scores (median RTs for low-probability triplets minus median RTs for high-probability triplets) by the *average* RTs of the given unit of five blocks for each participant and each unit of five blocks. The standardized scores were compared between groups with a mixed-design ANOVA with Block (Retrieval Phase 6-10 vs. Retrieval Phase 12-16 vs. Retrieval Phase 18-22) as a within-subject factor, and with Group (Dual Task Group vs. Single Task Group) as a between-subject factor. Please note that in the last two analyses, the dependent variable is a *difference score* of the high- and low-probability triplets (i.e., learning score).

For all ANOVAs, the Greenhouse-Geisser epsilon (ε) correction was used if necessary. Corrected *df* values and corrected *p* values are reported (if applicable) along with partial eta-squared (η_*p*_^2^) as the measure of effect size. LSD (Least Significant Difference) tests were used for pair-wise comparisons. In addition to the frequentist analyses, we conducted Bayesian ANOVAs or independent samples t-tests on the *learning scores* to overcome the limitations of null-hypothesis significance testing (Dienes, 2014) and gain statistical evidence for potential null-results. Bayes Factor (BF) reflects how well a model behaves compared to the null-model. In the case of BF_01_, the smaller the value is, the better the model predicts the data. The BF_01_ values of the null model are always 1 (containing the grand mean only) (Jarosz & Wiley, 2014).

In the case of the t-test, we report the BF_01_ value of the comparison. For the ANOVAs, we report the BF_01_ values of the best-fitting model, the BF_01_ factor of the best model containing the Group as a factor, as well as the change from prior to posterior inclusion odds of this factor (Wagenmakers et al., 2018). All of the frequentist analysis of the ASRT task was carried out by using IBM SPSS Statistics 25, and the Bayesian analyses were run in JASP (version 0.10, JASP Team, 2019).

## Results

### Did the two groups perform equally before the dual-task phase?

As expected, both groups showed statistical learning in the Learning Phase, and the degree of learning did not differ significantly between groups (see 2^nd^ paragraph of the Supplementary Materials). The level of consolidation proved to be similar in the two groups (see 3^rd^ and 4^th^ paragraph of the Supplementary Materials).

### Did the two groups differ in the test blocks?

First, we wanted to find out whether the two groups performed differently in the test blocks (when the Dual-Task Group performed the secondary task). A mixed-design ANOVA on the RT scores with Triplet (high-vs. low-probability) × Block (Retrieval Phase Blocks 6-10 vs. Retrieval Phase Blocks 12-16 vs. Retrieval Phase Blocks 18-22) × Group (Dual-Task Group vs. Single-Task Group) factors revealed a significant main effect of Block, *F*(1.665, 109.864) = 11.004, *p* < 0.001, η_*p*_^2^ = 0.143, highlighting that the average RTs accelerated during the Retrieval Phase. The significant main effect of Triplet, *F*(1, 66) = 254.192, *p* < 0.001, η_*p*_^2^ = 0.794, showed that the statistical knowledge was still detectable during the test blocks. Block × Triplet interaction was not significant, *F*(2, 132) = 2.259, *p* = 0.108, η_*p*_^2^ = 0.033, indicating that the degree of the statistical learning did not change significantly throughout the test blocks. The main effect of Group was significant, *F*(1, 66) = 13.107, *p* < 0.001, η_*p*_^2^ = 0.166, revealing that the average RTs were higher in the Dual-Task Group than in the Single-Task Group. It suggests that completing the secondary task during the ASRT gave rise to a general slowing down in the task. The Block × Group interaction was significant, *F*(2, 132) = 23.402, *p* < 0.001, η_*p*_^2^ = 0.262, indicating that the general RT decrease was only detectable in the Dual-Task Group (all blocks differed from each other, all *p* < 0.001); the Single Task Group did not show acceleration following the Learning Phase (neither block was different from the others: *p* > 0.086). Importantly, the Triplet × Group interaction did not reach significance, *F*(1, 66) = 0.521, *p* = 0.473, η_*p*_^2^ = 0.008, revealing that there was no statistically significant difference between groups in terms of the degree of statistical knowledge. This lack of difference did not change throughout the blocks, as revealed by a non-significant Block × Triplet × Group interaction, *F*(2, 132) = 0.211, *p* = 0.810, η_*p*_^2^ = 0.003, (Figure 2). The Bayesian mixed-design ANOVA (containing only the Block and Group factors, as it was carried out on the *learning scores*, see the Statistical analysis section) revealed that the best-fitting model is the null-model BF_01_ = 1. The best-fitting model that contained the Group as a factor incorporated the Group factor solely. The comparison of these two models revealed that the data are 3.408 times more likely under the null-model than the model that contains the Group factor. The amount of change from prior to posterior inclusion odds also speaks against the inclusion of this factor (BF_inclusion_ = 0.205). Therefore, the data suggests a similar level of statistical knowledge regardless of the block (Blocks 6-10 vs. Blocks 12-16 vs. Blocks 18-22) or the group (Single-Task Group or Dual-Task Group).

**Figure 2.**
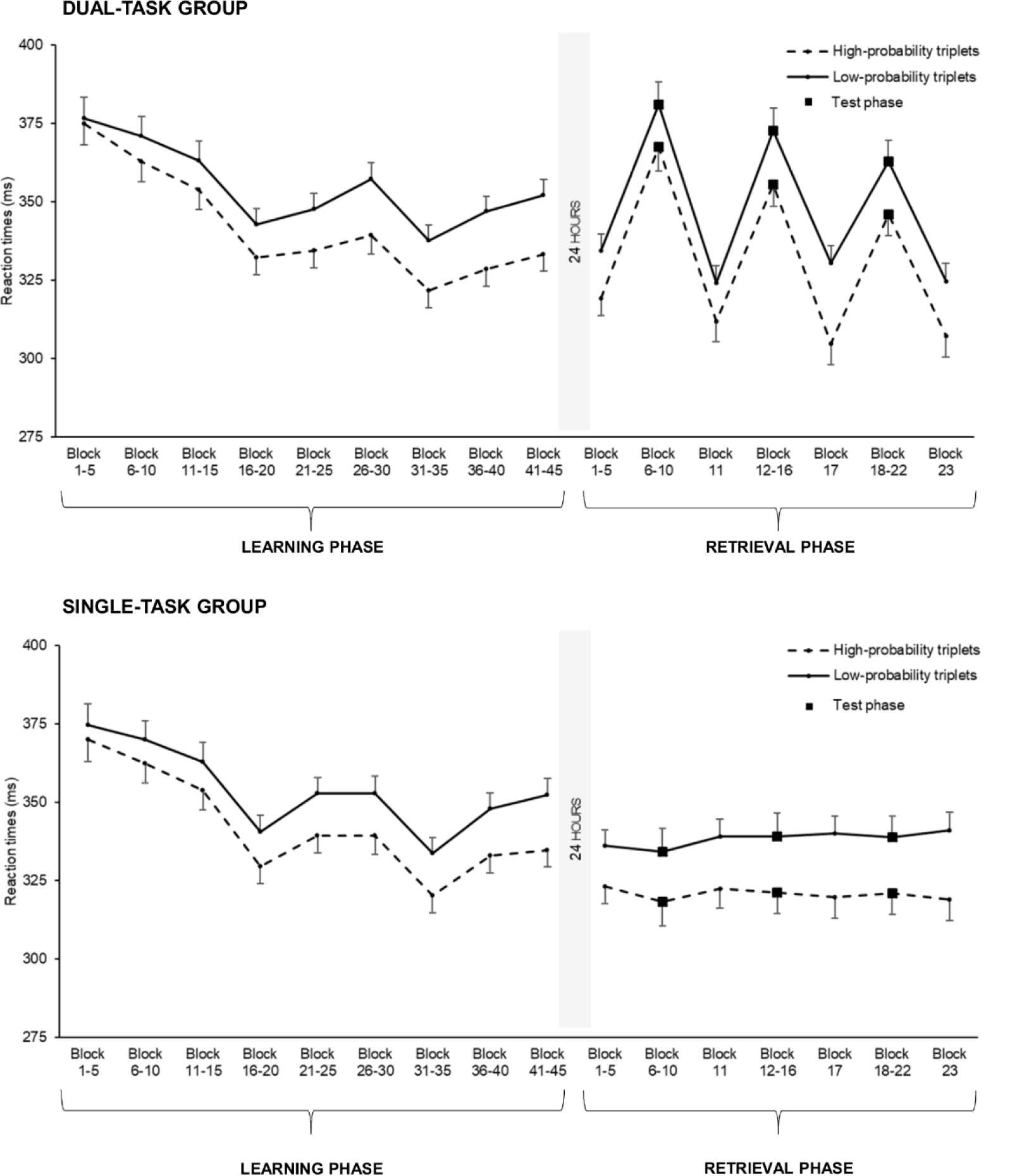
The RTs for the high- and low-probability triplets during the Learning and the Retrieval Phase, separatel for the two groups. On the vertical axes, we can see the median RTs in milliseconds, and on the horizontal axis, the nine units of five blocks of the Learning Phase (ST 1 – ST 45) and the four units of 5 blocks and the 3 probe blocks of the Retrieval Phase (Blocks 1-5, Blocks 6-10, Block 11, Blocks 12-16, Block 17, Blocks 18-22, Block 23). Th solid line represents the RTs for the low-probability triplets and the dashed line the RTs for the high-probabilit ones. The black squares represent the test phase when the Dual-Task Group performed the secondary task too. The error bars represent the standard error. The average RTs in the task became smaller over the Learning Phase, and the difference between the high- and low-probability triplets more pronounced. At the beginning of the Retrieval Phase, stable statistical knowledge was detected in Blocks 1-5. The dual-tasking slowed down the RTs of the participants, but the statistical knowledge remained stable even in these phases. We found similar results with standardized scores.

### Did the two groups differ in the probe blocks?

We checked if the groups performed differently in the probe blocks between the test blocks (Block 11, Block 17 and Block 23). As one block contains only 85 button presses, we averaged over the three blocks to gain more statistical power. The Triplet × Group ANOVA revealed a main effect of Triplet, *F*(1, 66) = 114.514, *p* < 0.001, η_*p*_^2^ = 0.634, revealing a stable statistical knowledge. The main effect of Group was not significant, *F*(1, 66) = 2.694, *p* = 0.105, η_*p*_^2^ = 0.039, indicating that the delaying effect of the dual-tasking did not significantly affect the performance on the probe blocks. Most importantly, the Triplet × Group interaction was not significant, *F*(1, 66) = 0.128, *p* = 0.721, η_*p*_^2^ = 0.002, revealing a lack of statistically significant difference between groups in terms of the degree of statistical knowledge also in the inserted probe blocks (Figure 3). The Bayesian independent samples t-test revealed substantial evidence for the lack of difference between groups (BF_01_ = 3.802).

**Figure 3.**
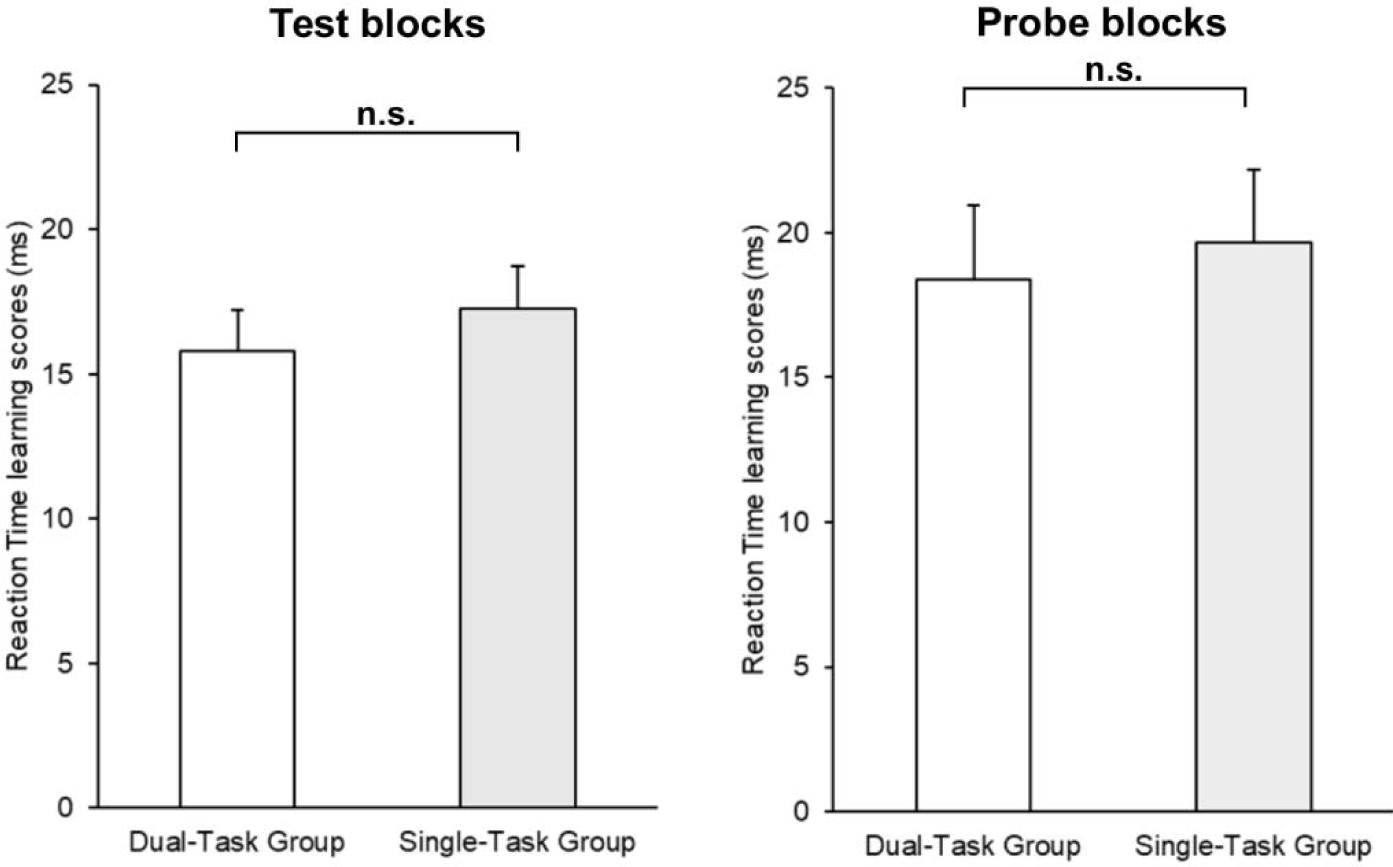
The RT learning scores of the target blocks and the probe blocks of the Retrieval Phase. The horizontal axis represents the two groups, and the vertical axis the RT learning scores (the RTs for the low-probability triplets *minus* the RTs for the high-probability triplets, the blocks collapsed together). The error bars signal the standard error. The learning scores of the two groups did not differ in the test blocks nor the probe blocks. We found similar results with standardized scores.

### Did the two groups differ in the test blocks in terms of standardized scores?

The Block × Group ANOVA of the *standardized RT learning scores* did not reveal a main effect of Block, *F*(2, 132) = 1.386, *p* = 0.254, η_*p*_^2^ = 0.021, suggesting that the learning scores did not change significantly during the test blocks. Importantly, consistent with the results without correction, no group difference was found in terms of the degree of statistical learning (Group: *F*(1, 66) = 1.325, *p* = 0.254, η_*p*_^2^ = 0.020). This lack of significant difference remained stable throughout the test blocks, as revealed by a non-significant Block × Group interaction, *F*(2, 132) = 0.018, *p* = 0.982, η_*p*_^2^ *<* 0.001. Similarly to the non-standardized results, the Bayesian mixed-design ANOVA suggested that the null-model is the best-fitting model (BF_01_ = 1). The best-fitting model that contains the Group as a factor involves the Group factor solely. However, the comparison of these two models revealed that the data are 2.508 times more likely under the null-model than the model that contains the Group factor solely. The amount of change from prior to posterior inclusion odds also speaks against the inclusion of this factor (BF_inclusion_ = 0.267). These results suggest a similar level of statistical knowledge regardless of the block (Blocks 6-10 vs. Blocks 12-16 vs. Blocks 18-22) or the group (Single-Task Group or Dual-task Group).

### How did the learning scores of the test blocks and the probe blocks compare?

We examined if the learning scores of the test blocks and the probe blocks differed from each other and whether it was similar between the two groups. The Block Type × Group ANOVA on the learning scores did not reveal the main effect of Block Type, *F*(1,66) = 1.872, *p* = 0.176, η_*p*_^2^ = 0.028, suggesting that there is no significant difference in the measured sequence knowledge between the test and probe blocks (i.e., between the periods where the stimulus stream contained colored stimuli). The main effect of Group did not reach significance, *F*(1,66) = 0.378, *p* = 0.541, η_*p*_^2^ = 0.006, indicating that the two groups performed similarly on the dual-task parts of the second session. More importantly, there was no statistically significant difference between the two groups in how the learning scores develop between the two types of blocks, as suggested by non-significant interaction of the Block Type and Group factors, *F*(1,66) = 0.004, *p* = 0.953, η_*p*_^2^ < 0.001. The Bayesian ANOVA revealed that the best-fitting model is the null-model (BF_01_ = 1). The best-fitting model that included the Group factor contained the Group factor solely, and the data proved to be 3.801 times more likely under the null-model than the model containing the Group factor that speaks against the inclusion of this factor (BF_inclusion_ = 0.191).

## Discussion

Here we investigated the effect of a secondary task on the retrieval of a well-established implicit sequence knowledge. Our participants practiced a sequence learning task with non-adjacent second-order dependencies through 45 blocks. After a 24-hour offline period, the participants were tested again on the task with or without a concurrent stimulus-counting task. Participants who were examined under dual-task conditions had retrieved their statistical knowledge to the same level as the participants with only single-task testing conditions. Moreover, this similarity proved to be true during blocks where both groups retrieved their knowledge under single-task conditions. These results remained the same, even when the differences in average RTs between groups was controlled. The lack of difference between groups in terms of the implicit sequence learning performance was also confirmed by Bayesian statistical methods. Our results went beyond previous literature by showing that well-learned, non-adjacent probabilistic sequence knowledge can be resistant to a concurrent secondary task.

Our study could have met three possible outcomes: impaired, intact, or improved retrieval of the learned probabilities under dual-task conditions compared to single-task retrieval. Expecting impaired performance can seem reasonable at first, as the majority of previous literature reported deteriorating effects of a secondary task on sequence learning abilities (Franklin et al., 2016; Hemond et al., 2010; Jiménez & Vázquez, 2005; Röttger et al., 2017; Shanks et al., 2005; Wierzchon et al., 2012). This deteriorating effect on learning was explained by numerous theories such as the suppression hypothesis (Frensch et al., 1998, 1999), task integration (Schmidtke & Heuer, 1997), organizational hypothesis (Stadler, 1995) or the response selection hypothesis (Schumacher & Schwarb, 2009). However, other studies did reveal intact or even improved learning, especially in the case of more complex statistical regularities (Jiménez & Méndez, 1999; Jiménez & Vázquez, 2005; Nemeth et al., 2011; Schvaneveldt & Gomez, 1998). An important difference between our and most previous studies is that we introduced the secondary task after a considerable amount of practice on the primary task. The participants completed more than 4000 trials on the primary task before the introduction of a dual-task, while the practice on the primary task ranged from zero to a few hundred trials in most of the previous studies. With this modification, we did not find evidence for impaired efficiency of the retrieval of the learned information under a dual-task condition. Contrary to our results, an early study of Schvaneveldt and Gomez (1998) had found impaired probabilistic sequence knowledge in a dual-task condition after initial single-task learning, but not on a single-task condition after initial dual-task learning. They had concluded that the information learned in single-task situations cannot be applied to a dual-task condition. However, in their study, they tested the transfer to dual-task conditions within one session (with less practice). On the contrary, we implemented a longer practice period, a 24-hour offline and a reactivation period to make sure that the sequence is well-learned before the retrieval. This suggests that acquiring the sequence knowledge to a great extent might help maintain a good level of retrieval during a subsequent dual-task condition.

Apart from the potentially disruptive effect of the secondary task, another possible outcome of the study was that the concurrent task would leave access to the sequence knowledge intact. This would mean that the processes behind the sequence knowledge retrieval and the stimulus-counting secondary task are independent from each other, similarly to how performance becomes automatized and resistant to dual-tasking on non-complex choice-response tasks (e.g. Brown & Bennett, 2002; Hazeltine et al., 2002; Logan, 1979; Ruthruff et al., 2001, 2006; Van Selst et al., 1999). As knowledge becomes skill-like, it ceases to rely on the same resources. Our main results are in line with this prospect: the degree of statistical knowledge remained the same compared to the single task retrieval during dual-tasking, as well as in the probe blocks, where both groups accessed their statistical knowledge without a concurrent counting task (please note that here the sequence knowledge is the one that becomes automatized but not the perceptual-motor improvement, see below). Moreover, the lack of differences persisted even after the normalization of the baseline reaction times. The fact that the sequence knowledge was comparable to the single-task group in both phases in the dual-task blocks (performance) and in the intermittent probe blocks (competence) indicates that the secondary task did not affect the performance nor the competence of the primary task (Kiss, Nemeth, & Janacsek, 2019; Vékony et al., 2019). These results are also in harmony with previous research that found intact implicit sequence knowledge after practicing the primary task in single-task conditions (Frensch et al., 1998; Shanks & Channon, 2002). However, in these studies, the presentation of the dual-task blocks followed immediately the few learning blocks, which makes it harder to draw conclusions about the retrieval after a long-term offline period. Therefore, our results extend this knowledge by providing evidence for three additional aspects. First, the retrieval of sequence knowledge remains resistant to a concurrent task even after a 24-hour offline period, which underlies the robust nature of probabilistic learning (Kóbor et al., 2017; Nemeth & Janacsek, 2011). Second, the retrieval of implicit probabilistic representations (see Supplementary Material) remains intact after extended practice. Third, neither the competence nor the performance of a well-acquired sequence knowledge can be disrupted by a secondary task.

The third potential outcome of the study was that the secondary task would improve the retrieval of the memory representations of the primary sequence learning task. This possibility would fit in well with the hypothesis of competition between control functions mediated by prefrontal cortical areas and sequence learning abilities. Several studies have discovered negative correlations between control functions and probabilistic sequence learning (Ambrus et al., 2019; Filoteo et al., 2010; Janacsek et al., 2012; Nemeth et al., 2013; Tóth et al., 2017; Virag et al., 2015), which suggests that a secondary task might potentially facilitate the access to sequence representations. Here we did not find improved performance in the dual-task condition as would be predicted by the competition theory. A possible explanation is that the competition theory is not applicable in dual-task situations. A more plausible explanation is related to the specificity and characteristics of the secondary task. Although the prolongation of reaction times during the secondary task confirms the effectiveness of our stimulus-counting task as a distraction, this task may not engage those specific mechanisms whose trigger the competitive interactions resulting in better performance in the retrieval of probabilistic sequence knowledge (Hemond et al., 2010). The exploration of which secondary tasks (if any) might be advantageous for sequence knowledge retrieval deserves future investigations.

Beyond the theoretical explanations, methodological aspects can also account for the results. The ASRT task allows us to disentangle the general skill-related processes and sequence-specific knowledge. The former was not taken into account by many previous studies, which makes it harder to unveil the underlying mechanism behind dual-task effects. In our study, the general skill-related processes such as perceptual-motor coordination and adaptation to the experimental situation are shown by the change of the overall RTs over time, while the sequence-specific knowledge is considered to be the emergence of difference between high and low-probability triplets (statistical learning). It is important to note that the increased overall RTs during the retrieval phase under dual-task conditions did not reveal impaired *sequence knowledge: T*hey indicate altered *general skill retrieval* on the primary task due to the dual-task constraint. The secondary task slowed down the overall perceptual-motor coordination, suggesting that in this aspect, the performance was not automatized until this point. However, the statistical knowledge that emerged during the learning phase became robust enough to persist under dual-tasking, thus we found a dissociation between the two processes. The dissection of the general skill learning and the statistical aspect of the learning process was also supported by the fact that in our data, after the normalization of the baseline RTs, the lack of differences in statistical learning between the single- and the dual-task group persisted. This result is crucial for stating that sequence knowledge was similar between the groups as general skill and sequence learning have been differentiated by previous studies (e.g. Juhasz, Nemeth, & Janacsek, 2019). Previous inconsistencies in the dual-task literature might also originate from differences in the proportion of general skill- and statistical learning-related factors of the used task. Therefore, future studies investigating the process of sequence learning or the retrieval of the learned knowledge under dual-task conditions could benefit from considering these aspects as potential confounding factors.

Previous studies have tried to determine which characteristics of the secondary tasks are crucial for disrupting the learning process, such as the correlation between the primary and the secondary task events or the features of the required response (e.g. Röttger et al., 2017). In our study, we chose a visual secondary task that is implemented in the stimuli stream, which does not break the stimulus-response interval therefore not causing interference as tone counting tasks might do (Jiménez & Méndez, 1999). However, we do not know if different secondary tasks involving functionally distant cognitive processes affect the sequence retrieval to a similar extent. For example, sentence processing has been found to impair probabilistic sequence learning, while mathematical and word processing tasks have not revealed disruptive effect (Nemeth et al., 2011). This result can be explained by the fact that language processing also relies on non-adjacent dependencies, similarly to the probabilistic sequence learning task used in the study. On the contrary, sequence learning has also been shown to be boosted when the secondary task involved similar sequence-learning mechanisms as the primary adjacent serial reaction time task (Hemond et al., 2010). Therefore, the set of cognitive processes that can and cannot interfere with the retrieval of sequence information is yet to be empirically established.

In everyday life, we mostly perform a secondary task when the primary one is well-acquired. In spite of this fact, no study has investigated the effect of a secondary task on the retrieval of well-acquired, complex statistical regularities. With the aim of filling this gap of knowledge, we exposed participants to a secondary task after extensive practice on the primary perceptual-motor sequence learning task. We found an intact retrieval of implicit probabilistic sequence knowledge, providing evidence that complex probabilistic representations can be robust against dual-tasking even if the general perceptual-motoric aspect of the primary task is affected. This suggests that complex statistical representations become more resistant to disruption than the general perceptual-motoric aspects and that we are able to correctly utilize our statistical knowledge if we are performing concurrently a new, non-statistical secondary task. Our results emphasize the importance of studying the dual-task effect not only in the learning phase but also during memory retrieval.

## Acknowledgments

This research was supported by the National Brain Research Program (project 2017-1.2.1-NKP-2017-00002); Hungarian Scientific Research Fund (NKFIH-OTKA K 128016, PI: D.N., NKFIH-OTKA PD 124148, PI: K.J.); Janos Bolyai Research Fellowship of the Hungarian Academy of Sciences (to K.J.); IDEXLYON Fellowship of the University of Lyon as part of the Programme Investissements d’Avenir (ANR-16-IDEX-0005) (to D.N). Thanks to Lison Fanuel for the comments and suggestions on the manuscript.

## Conflict of interest

The authors report no conflict of interest.

## Notes

### Competing Interest Statement

The authors have declared no competing interest.

