## Supplementary Materials for "Retrieval of a well-established skill is resistant to distraction: evidence from an implicit probabilistic sequence learning task"

**ASRT performance in the Learning Phase (matching criteria)**

***Analysis***

To make sure that the two groups performed equally before the dual-task phase, we compared the performance of the groups in the Learning Phase, between the two sessions and at the beginning of the second session. Reaction times were analyzed with mixed-designed ANOVAs with Triplet (high- vs. low-probability) and Blocks (Blocks 1-5 vs. Blocks 6-10 vs. Blocks 11-15 vs. Blocks 16-20 vs. Blocks 21-25 vs. Blocks 26-30 vs. Blocks 31-35 vs. Blocks 36-40 vs. Blocks 41-45) as within-subject factors and with Group (Dual-Task Group vs. Single-Task Group) as between-subject factor. To check whether the two groups consolidated the statistical knowledge to a similar extent between the two sessions, we compared the performances between the last five blocks of the first session (Learning Phase Block 41-45) of the Learning Phase and the first five blocks of the Retrieval Phase (Retrieval Phase Block 1-5) with a mixed-designed ANOVA with Triplet (high- vs. low-probability) and Blocks (Learning Phase Block 41-45 vs. Retrieval Phase Block 1-5) as within-subject factors and with Group (Dual-Task Group vs. Single-Task Group) between-subject factor. The performance at the beginning of the Retrieval Phase was also compared between groups with a mixed-designed ANOVA with Triplet (high- vs. low-probability) within-subject factor and with Group (Dual-Task Group vs. Single-Task Group) as between-subject factor.

***Did the two groups perform equally before the dual-task phase?***

First, we checked whether the two groups’ performance differed in the Learning Phase. The Block × Triplet × Group ANOVA of the RTs revealed a significant main effect of Block, *F*(3.922, 528) = 105.004, *p* < 0.001, η*_p_*^2^ = 0.614, revealing that participants generally become faster throughout the blocks. We found a significant main effect of Triplet, *F*(1, 66) = 325.545, *p* < 0.001, η*_p_*^2^ = 0.813, as RTs for high-probability triplets were smaller than for the low-probability ones, revealing statistical learning. Moreover, we found a significant Block × Triplet interaction, *F*(8, 528) = 44.735, *p* < 0.001, η*_p_*^2^ = 0.213, indicating that the degree of statistical learning increased throughout the blocks. We did not find the main effect of Group, *F*(1, 66) = 0.001, *p* = 0.966, η*_p_*^2^ < 0.001, suggesting that there was a statistically significant difference between groups in terms of average RTs. No group differences were found in the acceleration of average RTs during blocks (Block × Group: *F*(8, 528) = 0.935, *p* = 0.487, η*_p_*^2^ = 0.014), nor in the degree of statistical learning (Triplet × Group: *F*(1, 66) = 0.472, *p* = 0.494, η*_p_*^2^ = 0.007). The dynamics of the change in the degree of statistical learning also did not differ significantly between groups (Block × Triplet × Group: *F*(8, 528) = 0.926, *p* = 0.487, η*_p_*^2^ = 0.014).

After that, we checked whether there was a significant difference in the level of consolidation between groups after the 24-hour offline period. The Block × Triplet × Group ANOVA for the RTs revealed a significant main effect of Block, *F*(1, 66) = 78.493, *p* < 0.001, η*_p_*^2^ = 0.543, suggesting that participants generally become faster on the second day. Again, a significant main effect of Triplet was found, *F*(1, 66) = 364.855, *p* < 0.001, η*_p_*^2^ = 0.847, revealing statistical learning. The interaction of the Triplet and Block factors was significant, *F*(1,66) = 0.193, *p* = 0.005, η*_p_*^2^ = 0.115, indicating that the degree of statistical learning in RTs became smaller for the second session. We did not find statistically significant difference between groups in terms of average RTs, *F*(1, 66) = 0.061, *p* = 0.806, η*_p_*^2^ = 0.001. No group differences emerged in the general RTs change between blocks (Block × Group: *F*(1, 66) = 0.302, *p* = 0.584, η*_p_*^2^ = 0.005), nor in the degree of statistical learning (Triplet × Group: *F*(1, 66) = 1.042, *p* = 0.311, η*_p_*^2^ = 0.016). The dynamics of the change in the degree of statistical learning between blocks did not differ significantly between groups (Block × Triplet × Group: *F*(1, 66) = 0.193, *p* = 0.662, η*_p_*^2^ = 0.003).

Finally, we compared the performance in the five blocks of the Retrieval Phase (ST 1-5), to check whether there was a difference between groups before the dual-task phase. The Triplet × Group ANOVA showed a significant main effect of Triplet, *F*(1, 66) = 195.812, *p* < 0.001, η*_p_*^2^ = 0.748, revealing again statistical learning effect. However, no statistically significant difference between groups was found neither in terms of average RT (Group: *F*(1, 66) = 0.130, *p* = 0.719, η*_p_*^2^ = 0.002), nor in the degree of statistical learning (Triplet × Group: *F*(1, 66) = 1.349, *p* = 0.250, η*_p_*^2^ = 0.020).

**Inclusion-Exclusion Task**

We assessed the “*Process Dissociation Procedure (PDP)*” (Jacoby, 1991) by administering the Inclusion-Exclusion Task (Destrebecqz & Cleeremans, 2001; Destrebecqz et al., 2005; Fu, Dienes, & Fu, 2010; Jiménez, Vaquero, & Lupiáñez, 2006). Employing this task, we revealed whether the participants gained explicit conscious knowledge about the statistical regularities during the ASRT task. Before administering the task (after the two sessions of ASRT), we informed the participants that the appearance of the stimuli followed a regularity. Then, in the first part of the task, we asked them to generate a sequence of button presses that follows the regularity of the ASRT task, using the same four response buttons the participants used during the ASRT task (*Inclusion condition*). They performed four runs of the Inclusion condition, and each run finished after 24 button presses, which is equal to three rounds of the eight-element alternating sequence (Horvath, Torok, Pesthy, Nemeth, & Janacsek, 2018; Kiss, Nemeth, & Janacsek, 2019; Kobor, Janacsek, Takacs, & Nemeth, 2017). After that, participants were asked to generate new sequences of responses that are *different* from the learned one (*Exclusion condition*). The Exclusion condition also contained four runs.

According to the PDP, participants can achieve successful performance in the Inclusion condition by solely implicit knowledge (explicit knowledge can also boost performance, but it is not necessary to the successful completion of the task). However, successful performance (i.e., intentionally generating *different* sequences) in the exclusion condition can only occur if the participant has conscious knowledge about the learned statistical regularities (if the participant knows what to suppress). Generation of the learned statistical regularities above chance level even in the Exclusion task indicates that the participants rely on their implicit knowledge, as they cannot control the generation of the learned sequences consciously. To test whether the participants gained consciously accessible triplet knowledge, we calculated the percentage of producing high-probability triplets in the Inclusion and the Exclusion condition separately. We tested whether it differs from the probability of generating them by chance. We also compared the percentages of the high-probability triplets across conditions (Inclusion and Exclusion task) and groups (Dual-Task Group and Single-Task Group) (for more details about the Inclusion-Exclusion task, see: Horvath et al., 2018; Kiss et al., 2019; Kobor et al., 2017).

***Analysis***

To check whether the participants developed conscious knowledge about the learned probabilistic regularities, we compared the probability of high-probability triplets in the generated sequences in the Inclusion-Exclusion test to chance level (25%) with the help of one-samples t-tests, separately for the two conditions (Inclusion and Exclusion). We also compared the performance in the Inclusion and in the Exclusion condition separately for the two groups with paired-sampled t-tests. Finally, we compared the probability of the high-frequency triplets between the two groups with independent samples t-test for the Inclusion and for the Exclusion condition as well. The statistical analysis was also carried out by using IBM SPSS Statistics 25.

***Did the participants develop conscious knowledge about the statistical regularities and was it different between groups?***

To test whether the acquired statistical knowledge remained implicit or became intentionally accessible, the Inclusion/Exclusion task was administered (see Methods for details). In the Dual-Task Group, 5 participants were excluded from this analysis, as they were apparently not following the instruction properly in the Exclusion condition (they did not generate diverse sequences). The participants of the Dual-Task Group generated 6.03% more high-probability triplets than chance level (25%), *t*(28) = 4.072, *p* < 0.001, in the Inclusion condition, and 4.45% more high-probability triplets in the Exclusion condition, *t*(28) = 2.308, *p* = 0.029. Comparing the two conditions, no significant difference was found, *t*(28) = 0.807, *p* = 0.426. In the Single-Task Group, 7 participants were excluded in total, as 6 participants did not follow the instructions in the Exclusion condition and 1 participant in the Inclusion condition. Overall, participants in this group generated 5.86% more high-probability triplets in the Inclusion condition than chance level, *t*(26) = 5.530, *p* < 0.001, and 4.67% more in the Exclusion condition, *t*(26) = 2.412, *p* = 0.023. Comparing the two conditions, no significant difference was found, *t*(26) = 0.597, *p* = 0.556. Comparing the performances of the two groups, no differences were found neither in the Inclusion, *t*(54) = 0.092, *p* = 0.927, nor in the Exclusion condition, *t*(54) = -0.078, *p* = 0.938. Taken together, the results suggest that both groups acquired the triplet knowledge, but could not consciously access and control it.
